# A universal preservation protocol for multi-omic and histological analysis of kidney tissue

**DOI:** 10.1101/2023.08.16.553482

**Authors:** Sydney E. Gies, Sonja Hänzelmann, Dominik Kylies, Simon Lagies, Moritz Lassé, Fabian Hausmann, Robin Khatri, Manuela Poets, Tianran Zhang, Shun Lu, Shuya Liu, Silvia Chilla, Ilka Edenhofer, Jan Czogalla, Fabian Braun, Bernd Kammerer, Markus M. Rinschen, Victor G. Puelles, Stefan Bonn, Maja T. Lindenmeyer, Tobias B. Huber

## Abstract

Biobanking of tissue from clinically obtained kidney biopsies for later use with multi-omic and imaging techniques is an inevitable step to overcome the need of disease model systems and towards translational medicine. Hence, collection protocols ensuring integration into daily clinical routines using preservation media not requiring liquid nitrogen but instantly preserving kidney tissue for clinical and scientific analyses are of paramount importance. Thus, we modified a robust single nucleus dissociation protocol for kidney tissue stored snap frozen or in the preservation media RNA*later* and CellCover. Using porcine kidney tissue as surrogate for human kidney tissue, we conducted single nucleus RNA sequencing with the Chromium 10X Genomics platform. The resulting data sets from each storage condition were analyzed to identify any potential variations in transcriptomic profiles. Furthermore, we assessed the suitability of the preservation media for additional analysis techniques (proteomics, metabolomics) and the preservation of tissue architecture for histopathological examination including immunofluorescence staining. In this study, we show that in daily clinical routines the RNA*later* facilitates the collection of highly preserved kidney biopsies and enables further analysis with cutting-edge techniques like single nucleus RNA sequencing, proteomics, and histopathological evaluation. Only metabolome analysis is currently restricted to snap frozen tissue. This work will contribute to build tissue biobanks with well-defined cohorts of the respective kidney disease that can be deeply molecularly characterized, opening new horizons for the identification of unique cells, pathways and biomarkers for the prevention, early identification, and targeted therapy of kidney diseases.

## Introduction

Independent of the underlying etiology, chronic kidney disease (CKD) constitutes a major risk factor for increased morbidity and mortality as well as progression to end stage kidney disease. Approximately 10% of adults worldwide are affected by CKD and the global burden of chronic kidney disease is growing: CKD is estimated to become the fifth leading cause of death globally by 2040 [1]. In recent decades, scientific effort has shed light on the various pathomechanisms of kidney diseases and the resulting therapies, including renal replacement by dialysis or transplantation. However, most findings are based on model systems (animal and cell models) of the respective kidney disease, limiting the options for targeted therapy and identifying prognostic biomarkers. Therefore, access to high-quality human tissue samples is an important pillar of translational research. The sample quality itself is influenced by various factors such as sample collection, storage, and processing. Biobanks represent an excellent platform for collecting kidney needle biopsies in relevant numbers under standardized conditions, even for rare diseases, and for building well-defined cohorts.

A major challenge for clinically obtained biopsies is the workflow integration into everyday clinical routine, as snap freezing still represents the gold standard for cryopreservation of kidney tissue for certain applications. Human kidney biopsies collected for clinical diagnostics are obtained during the day and often geographically distant from the laboratory, impeding immediate transfer and processing of fresh tissue and snap freezing in liquid nitrogen. A common approach to circumvent these obstacles is to harvest tissue samples for biobanking in cryopreservative agents, such as RNA*later* [2], [3], [4], [5]. RNA*later* is a commercially available, aqueous, nontoxic ammonium sulfate containing tissue storage reagent that preserves instantly the integrity of cellular RNA in tissue samples even at room temperature for up to one week and has shown to be sufficient for long-term protection of cellular RNA at - 20°C or −80°C, thus minimizing the need of rapid processing or freezing tissue samples in liquid nitrogen [6], [7], [8]. Recent publications demonstrate that single nucleus dissociation protocols applied to solid tissues such as the kidney stored in RNA*later* at −80°C were suitable for further analyses with single nucleus (sn) RNA sequencing (snRNAseq) protocol [9]. However, to the best of our knowledge, neither the possible effects of RNA*later* on the sn transcriptome nor the suitability of RNA*later* stored kidney tissue for proteomics, metabolomics or preservation of kidney tissue architecture and immunofluorescence staining has been examined.

The newer formulation CellCover protects DNA, RNA and proteins of cells and solid tissues by instant disruption of cellular degradation pathways immediately after exposure without applying low temperatures. No data exists about storage of CellCover preserved kidney tissue at −80°C for subsequent multi-omic approaches.

Single cell (sc) RNAseq was demonstrated to identify unique cells, pathway activity, consecutive targets for novel therapies and preventive and prognostic markers [10], [11], [12], thus supplementing or supporting data derived with other techniques such as bulk RNAseq, proteomics, metabolomics as well as the multiple, complementary types of histological staining. Therefore, a standard for cryopreserving kidney tissue for later deep molecular characterization with multi-omic approaches focusing on snRNseq even months or years after collection is an inevitable step to ensure comparability and reproducibility of data.

As human kidney tissue is very valuable and limited, we have used pig kidneys as a surrogate tissue for the studies. Many protocols are optimized for mouse tissue, which is not comparable to healthy and diseased human kidney tissue in terms of fibrotic portions. Pig kidneys, on the other hand, are anatomically very similar to human kidneys, can be easily harvested at local slaughterhouses and no application for animal testing permits is required. Thus, they represent an innovative approach for the development of protocols to reduce the use of laboratory animals, which aligns with the 3Rs principles (replacement, reduction, refinement) in animal research and conserve human tissue.

## Materials and Methods

### Pig kidney tissue collection and cryopreservation

Fresh pig kidneys were obtained from a local slaughterhouse and immediately placed on ice. To mimic clinical settings, the pig kidneys were biopsied with a 16G biopsy needle. The biopsy cores were either immediately frozen in liquid nitrogen or put into 1 ml of RNA*later* (AM7020, Invitrogen) or CellCover (800-050, Anacyte) for 30-60 minutes at 4°C. After incubation, the pig kidney biopsies in RNA*later* and CellCover were placed into a −80°C freezer for at least one month.

For immunohistochemical analysis, different freezing conditions were applied to pig kidney biopsies after they incubated for 30-60 minutes in RNA*later* or CellCover at 4°C. They were frozen to −20°C with or without using a CoolCell (479-1840, Corning) and half of the samples were put at −80°C after 7 days. As controls, pig kidney biopsies were embedded in cryomolds (4557, Sakura) with optimal cutting temperature (OCT) medium (4583, Sakura) and frozen in liquid nitrogen. For staining, RNA*later* stored samples were thawed, rinsed with phosphate buffered saline (PBS), formalin-fixed and paraffin-embedded. CellCover stored kidney tissue was thawed and directly paraffin embedded.

### Human kidney tissue

For generating snRNAseq data of human kidney tissue, the second core of a clinical indicated allograft needle biopsy and 2 tumor-nephrectomy samples were put in 1ml of RNA*later* for around 60 minutes at 4°C and finally stored at −80°C. Written informed consent was obtained from all individuals and the local ethics committee approved of the studies (PV5026 and PV5822).

### Kidney tissue dissociation, library preparation and sequencing

#### Single nucleus dissociation

A kidney biopsy core was thawed on ice and 3-4 mm of the core were chopped with a razor blade in a petri dish on ice and homogenized using a Dounce homogenizer (D8938-1 SET, Sigma-Aldrich) in 200 µl ice-cold lysis solution and incubated on ice for 20 min with additional 3.8 ml of ice-cold lysis solution. If the biopsy core was larger as the 3-4 mm used for single nucleus dissociation, it was put back in RNA*later* and frozen again at −80°C. Lysis solution was prepared with Nuclei PURE lysis buffer (NUC-201, Sigma-Aldrich), 1 mM dithiothreitol (D9779, Sigma-Aldrich) and 0.1 % Triton X-100 (NUC-201, Sigma-Aldrich) according to manufacturer protocol and a RNAse inhibitor mix (0.04 U/µl SUPERaseIN RNAse Inhibitor [AM 2696, Thermo Fisher]; 0.04 U/µl RNAsin Plus RNAse Inhibitor [N2615, Promega]) was added. The single nuclei suspension was filtered through a 30 µm strainer (04-004-2326, Sysmex) and centrifuged at 500g for 5 min at 4°C. The pellet was resuspended and incubated for 2 min in 1ml red blood cell lysis buffer hybri-max™ (R7757-100ml, Sigma Aldrich), filtered through a 5 µm strainer (04-004-2323, Sysmex) and washed with 4 ml of ice cold 0,01% BSA (AM2616, Thermo Fisher) in DPBS (59331C; Sigma) with 0.04 U/µl SUPERaseIN RNAse Inhibitor and 0.04 U/µl RNAsin Plus RNAse Inhibitor at 500 g for 5 min at 4°C. The pellet was resuspended in 1% BSA in DPBS with 0.04 U/µl SUPERaseIN RNAse Inhibitor and 0.04 U/µl RNAsin Plus RNAse Inhibitor.

#### Nuclei loading to the Chromium 10X platform

Nuclei number was counted and samples were diluted prior to loading to the Chromium 10X device to capture ∼ 6 000 nuclei of one snap frozen, one RNA*later* and the CellCover stored sample and to capture ∼ 10 000 nuclei of two snap frozen and one RNA*later* stored sample The nuclei were separated into Gel Bead Emulsion droplets and libraries were prepared with the Chromium NEXT GEM Single Cell 3’ Reagent kits v3.1 according to manufacturer’s protocol. The libraries were sequenced on an Illumina Novaseq6000 platform as symmetric paired end run (150 bases) with 200 million raw sequencing reads per sample (pig) or with 300 million raw sequencing reads per sample (human). Raw data and count tables for pig can be downloaded from GEO (GSE239442) and for human at Forschungsdatenrepository Hamburg (10.25592/uhhfdm.13026).

### Bioinformatics analyses

#### Preprocessing and QC

10x Genomics raw sequencing data were aligned using the CellRanger software suite (version 6.1.1, 10x Genomics, Pleasanton, CA) including intronic counts. The pig genome Sscrofa11.1 (GCA_000003025.6) was built following the instructions of the 10x Genomics web page for custom genomes using the GTF file (https://www.ensembl.org). The human genome (GRCh38-2020-A) was downloaded from 10x Genomics. Next, SoupX [13] (v1.6.1) was applied with default parameters to remove ambient RNA contamination. Filtering was done by removing cells with < 400 and > 5000 expressed genes. Furthermore, cells with more than 5% mitochondrial expression were excluded. Doublets were removed using Scrublet [14] (default parameters, v0.2.2). Genes, which were expressed in less than 3 cells, were removed.

#### Dimensionality reduction and clustering

After obtaining the sequencing data gene count matrix, subsequent quality control, normalization steps and correction of batch effects (Harmony [15]) provides the basis for a robust downstream analysis. The gene-cell-barcode matrix was log-transformed. PCA was run on the normalized gene-barcode matrix of the top 2000 most variable genes to reduce the number of feature (gene) dimensions. We used Scanpy [16] to perform dimensionality reduction with Uniform Manifold Approximation and Projection (UMAP) to visualize the data and identification of cell populations in combination with the Leiden clustering algorithm (resolution=0.31). The first 20 PCs and 30 neighbors were used for UMAP visualization.

#### Cell type identification based on marker genes and their expression

To identify genes that are enriched in a specific cluster, the mean expression of each gene was calculated across all cells in the cluster. Then each gene from the cluster was compared to the median expression of the same gene from cells in all other clusters. For each cluster, the marker gene list was determined by adjusted p value < 0.05 (sc.rank_genes_groups) and log2FoldChange > 0.5. The highly ranked genes were then used to determine the cell type of each cluster.

#### Proteomic analysis of kidney biopsies

Samples were prepared as previously described with small modifications [17]. In brief, 1-2 mm^3^ sections of biopsies were washed three times in PBS and then lysed in 8 M urea, 50 mM ammonium bicarbonate, supplemented with 1x Halt protease inhibitor cocktail (Thermo Scientific). Samples were homogenized using glass Dounce homogenizers and then sonicated. Proteins were then precipitated using ice cold acetone to remove any potentially interfering detergents. Protein pellets were re-solubilized in 8M urea, 50 mM ammonium bicarbonate, reduced with 5 mM DTT and alkylated using 10 mM IAA. Overnight digestion was carried out using trypsin at 1:50 enzyme to substrate ratio at 37 °C. Tryptic peptides were acidified to pH 3 using formic acid and purified using in-house made stage-tips [18]. 1 µg of protein was used for LC-MS/MS acquisition.

Proteomics data acquisition was carried out on a quadrupole Orbitrap mass spectrometer (QExactive; Thermo Fisher Scientific, Bremen, Germany) coupled to a nano UPLC (nanoAcquity system, Waters) with an inline trap column for desalting and purification (180 µm × 20 mm, 100 Å pore size, 5 µm particle size, Symmetry C18, Waters) followed by a 25 cm C18 reversed-phase column for peptide separation (75 µm × 200 mm, 130 Å pore size, 1.7 µm particle size, Peptide BEH C18, Waters). Peptides were separated using an 80-minute gradient with linearly increasing ACN concentration from 2% to 30% ACN in 65 minutes using a two-buffer system (buffer A: 0.1% FA in water, buffer B: 0.1% FA in ACN). The mass spectrometer was operated in data-dependent acquisition (DDA) mode with the top 15 ions by intensity per precursor scan (1 × 10^6^ ions, 70,000 Resolution, 240 ms fill time) being selected for MS/MS (HCD at 25 normalized collision energy, 1 × 10^5^ ions, 17,500 Resolution, 50 ms fill time) in a range of 400-1200 m/z. A dynamic precursor exclusion of 20 seconds was used. LC-MS/MS data were searched against the UniProt pig reference proteome (downloaded June 2021, 22166 entries) using MaxQuant (version 1.6.3.4) with default parameters. The match between runs (MBR) was disabled, LFQ, IBAQ and classical normalization features were enabled. Data analysis was performed using Perseus software suite (v1.6.15.0) [19]. Missing data was accepted to a threshold of 33% across samples, and missing data were imputed sample wise with a width of 0.3 SD and a downshift of 1.8 SD. Differences were assessed by ANOVA followed by Tukey’s HSD.

The mass spectrometry proteomics data including FASTA sequences for database searching have been deposited to the ProteomeXchange Consortium via the PRIDE [20] partner repository with the data set identifier PXD044127.

#### Metabolome analysis of cryopreserved pig kidney tissue

Kidney biopsies were grinded in liquid nitrogen before metabolites were extracted as previously [21] described. In brief, metabolites and lipids were simultaneously extracted by adding 1 ml methanol:water (1:1, including internal standards for normalization) and 0.5 ml chloroform (containing internal standard for normalization). The upper phase was subjected to non-targeted metabolomics by gas-chromatography coupled to mass spectrometry (GC-MS), while the lower phase was analyzed by targeted lipid profiling with liquid-chromatography (LC) coupled to multiple-reaction-monitoring (MRM) mass spectrometry. Details about chromatography, mass spectrometry and MRM-transitions can be found in Lagies et al. [21]. Data were log-transformed, statistical analysis and visualization were performed by MetaboAnalyst 5.0 [22].

### Immunofluorescence staining of cryopreserved pig kidney tissue

#### De-waxing and antigen retrieval

Paraffin-sections were cut at a thickness of 2-4µm and mounted on SuperFrost plus slides. Then, all samples were sequentially immersed in xylene 3x (10 minutes each) followed by an ethanol series (5 minutes each) of 3x 100%, 2x 70%, 1x 50% and finally 3x (5 minutes each) in double deionized water. Antigen retrieval was performed with Agilent DAKO Target Retrieval Solution pH9 (S236884-2) in a Braun Multiquick FS20 steamer for 15 minutes, followed by a cool-down to room temperature of 30 minutes. The sections were then incubated in Agilent Wash Buffer Solution (K800721-2) for 15 minutes at room temperature.

#### Fluorescent immunolabeling

Samples were incubated with primary antibodies at concentrations according to vendor’s guidelines in Agilent Antibody Diluent Solution (CK800621-2) overnight at 4°C, followed by three times washing for 5 minutes with Agilent Wash Buffer Solution. Then, sections were incubated with appropriate secondary antibodies as well as directly conjugated primary antibodies at concentrations according to vendor’s guidelines with DAPI (Sigma Aldrich D9542) at a final concentration of 1μg/ml in Agilent Antibody Diluent Solution for 1 hour at room temperature and washed again three times for 5 minutes with Agilent Wash Buffer Solution. All primary antibodies used in this publication are listed in Supplementary Table 1. The following secondary antibodies were used: goat anti-guinea pig Alexa Fluor 555 (Invitrogen A21435), goat anti-rabbit Alexa Fluor 647 (Invitrogen A21245). After immunostaining, samples were mounted with ProLong Gold (Invitrogen P36930).

#### Fluorescence imaging of tissues

LED-based widefield imaging of tissues was performed using the THUNDER Imager 3D Live Cell and 3D Cell Culture (Leica Microsystems). Low magnification images were obtained in combination with a 20x objective (NA: 0.40). High magnification images were obtained in combination with a 40x objective (NA: 1.10) after optimizing LED-intensity and exposure times. Fiji imaging software (Max Planck Institute of Molecular Cell Biology and Genetics) was used to navigate the raw files.

## Results

### Time and tissue saving single nuclei dissociation with standardized lysis buffer

snRNAseq has shown advantages over scRNAseq regarding activation of gene expression artifacts due to warm single cell dissociation protocols, viability, especially of cryopreserved tissue and proportionate cell-type recovery of all major cell types represented in the kidney [23]; [24], [10]. Therefore, we first optimized a single nuclei isolation protocol using a commercially available and thus standardized lysis buffer for cryopreserved kidney tissue that was either snap frozen, stored in RNA*later* or CellCover at −80°C. The pilot studies were performed on needle biopsies of pig kidneys. Furthermore, we reduced filtering and centrifugation steps and performed the entire procedure on ice to minimize gene expression artifacts, as at 4°C the mammalian transcriptional machinery is largely inactive [25]. Cryopreservation in RNA*later* and CellCover showed sufficient single nucleus dissociation of pig kidney biopsies after only 25 minutes, while snap frozen pig kidney needed 20 minutes more for a proper dissociation, as examination of nuclei morphology by light microscopy demonstrated (Supplementary Figure 1 A and B). The efficacy of the dissociation protocol was not influenced by the different cryopreservation methods. 3-4 mm of a thawed biopsy core resulted in 300 000-400 000 nuclei independent of the storage condition (data not shown) thus creating the possibility to complement single nucleus sequencing data with data obtained from other analyzing techniques of the same biopsy core. The procedure from thawing the biopsy core to counting and diluting the single nuclei suspension can be completed in approximately 60 minutes for immediate processing with any single cell platform.

### SnRNAseq of cryopreserved pig kidney tissue

Isolated nuclei from each storage condition were further processed with the Chromium 10X platform. Agilent Bioanalyzer traces of cDNA after reverse transcription/amplification and library construction showed the expected fragment size for each cryopreservation condition (Supplemental Figure 1 C, D and E).

After sequencing and processing of the Chromium 10X data sets with CellRanger software, the estimated number of cells was on average 9530 nuclei per sample, that was near to the targeted recovery of 10 000. On average, 26 676 total genes and a median number of 2421 genes per nucleus were detected. Q30 scores of UMIs, RNA reads and barcodes were mostly above 91.5% (Supplemental Table 2). Counts were normalized such that the total count of each cell was 10,000. The number of detected genes strongly correlated with the total number of RNA reads for snap frozen, RNA*later* and CellCover stored (R^2^= 0.94, 0.95, 0.94, Figure 1 A). Significant differences in the number of detected genes and the depth of gene expression were observed between RNA*later* and CellCover stored kidney tissue compared to snap frozen kidney tissue (p< 0.05, Figure 1 B and C). The total number of RNA reads showed no or only weak correlation with the Mitochondria-associated gene expression for snap frozen, RNA*later* and CellCover preserved samples (R²=0.15, 0.27 and 0.25, respectively, Figure 2A). However, RNA*later* and CellCover stored samples showed significantly higher fractions of mitochondrial counts (p< 0.05, Figure 2B). Besides mitochondrial genes, ambient RNA released during tissue-dissociation can contaminate snRNAseq data sets. It was reported that the highly expressed genes especially from highly abundant cells like the proximal tubule were detected in all cell clusters [11], [23]. Lowest ambient RNA contamination with a fraction of 0.055 of all counts was found in RNA*later* compared to snap frozen and CellCover stored kidney tissue (0.066 and 0.086 of all counts, Figure 2C). Nevertheless, the removal of the ambient RNA led to a significant reduction of the number of counts in each condition (Figure 2D).

**Fig. 1:**
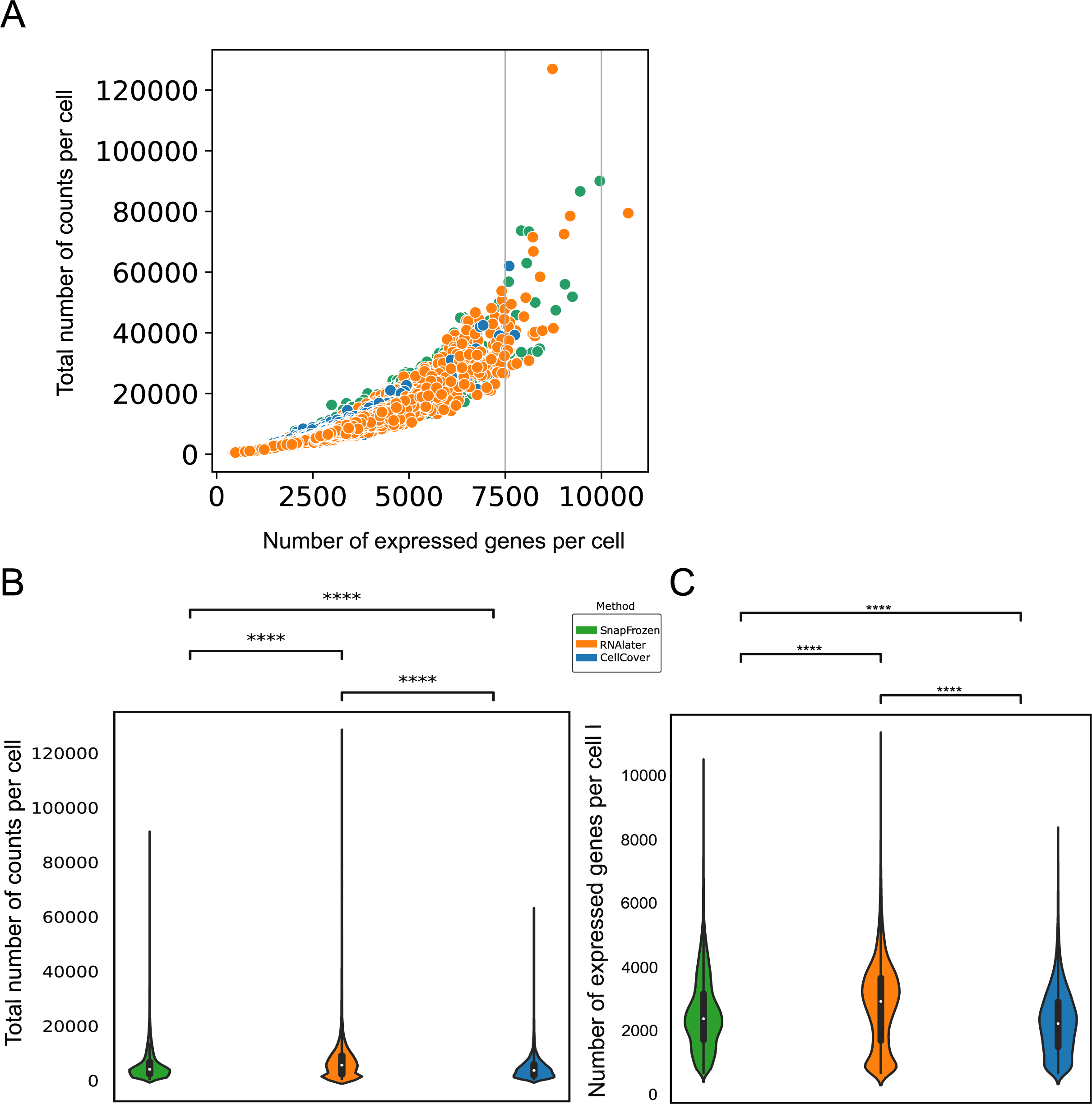
Scatter plot representation depicting the relationship of the number of detected genes (y-axis) against the total number of RNA reads (x-axis) for snap frozen (R^2= 0.94), RNA*later* (R^20= 0.94) and CellCover (R^2=0.95) stored kidney tissue (A). Comparison of the sequencing quality metrics total number of RNA reads (B) and total number of genes (C) between snRNAseq data sets obtained from different cryopreservation methods (snap frozen, RNAlater and CellCover stored kidney tissue). ****= p < 0.05

**Fig. 2.:**
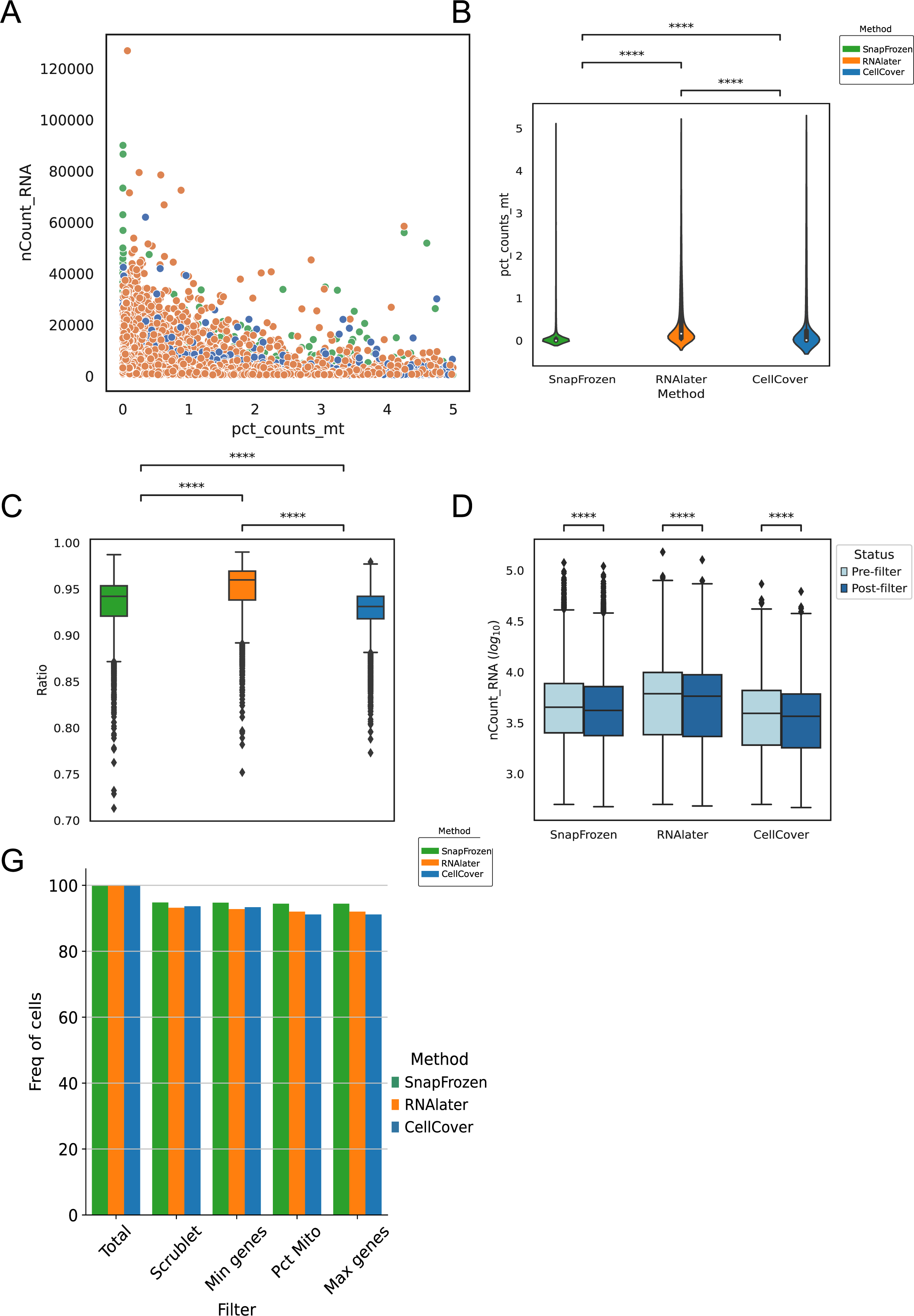
Comparison of quality metrics of 6 snRNAseq data sets from 3 snap frozen, 2 RNAlater and 1 CellCover cryopreserved pig kidney biopsies. (A) Correlation of the total number of molecules detected within a nucleus with percentage of mitochondria present in each cell averaged for each method. (B) Distribution of percentage of mitochondria in each cell per method. (C) Distribution of the ratio of ambient RNA and total RNA as reported by SoupX. (D) Distributions of nuclei containing ambient RNA and after removal with SoupX. The counts are log transformed for better visualization. (E) Number of remaining nuclei after each filtering step, including scrublet for multiplet removal, removal of nuclei with < 400 genes (min genes), removal of nuclei with > 5% mitochondrial counts (pct mito) and removal of nuclei with > 5000 genes (max genes). ****= p<0.05

After filtering was performed, the recovered number of nuclei was assessed for each storage condition to examine an impact of the storage condition on snRNAseq data quality. Over 90% of the nuclei of data sets from snap frozen, RNA*later* as well as CellCover stored kidney passed the filtering steps and no significant differences between the different cryopreservation methods were observed (Figure 2E). After applying all filtering steps to snRNAseq data sets, 31185, 15424 and 6742 nuclei with a median number of 2104, 2645 and 1949 genes/nucleus were recovered in snRNAseq data sets from three snap frozen, two RNA*later* and one CellCover sample stored kidney tissue, respectively.

### Assessment of preservation-related artifacts in snRNAseq data sets

The quality of the snRNAseq data sets was further evaluated by the ability to define cell types after dimensionality reduction in combination with clustering algorithm (see methods “dimensionality reduction and clustering” for a more detailed description). The six data sets were combined using Harmony [15]. The final combined data set included 53351 nuclei. UMAP analysis revealed that each sample processed with the described protocol contributed to 31 clusters (Figure 3A-C). These clusters showed unique marker gene expression of both common and rare kidney cell types (Fig. 3D-G), including podocytes (POD), 7 populations of endothelial cells (EC, Figure 3F) including glomerular (EC-1, gEC) and lymphatic endothelial cells (EC-7), mesangial cells (MC), fibroblasts (FIB), myofibroblasts (MyoFIB), vascular smooth muscle cells/pericytes (vSMC/P), vascular smooth muscle cells/mesangial cells (vSMC/MC), juxtaglomerular cells (JUXT), interstitial cells (INT), 3 populations of proximal tubules (PT, Figure 3E), tubule cells with markers of proximal tubule cells and podocytes (PODO-PT), thick ascending limb (TAL), distal convoluted tubules (DCT), tubule cells with markers of distal convoluted and connecting tubule cells (DCT/CNT), principal cells (PC), intercalated cells type A and B (IC-A, IC-B), 2 populations of monocytes/macrophages (MAC), t-cells (T-CELL, Figure 3G). No cluster heterogeneity was observed between the different storage conditions (Fig. 3A and C). Proximal tubules showed the highest frequency of all cell types under each cryopreservation condition and the cryopreservation method did not hinder the detection of cells that are present in low proportions in the kidney (Fig.3A and C).

**Fig. 3:**
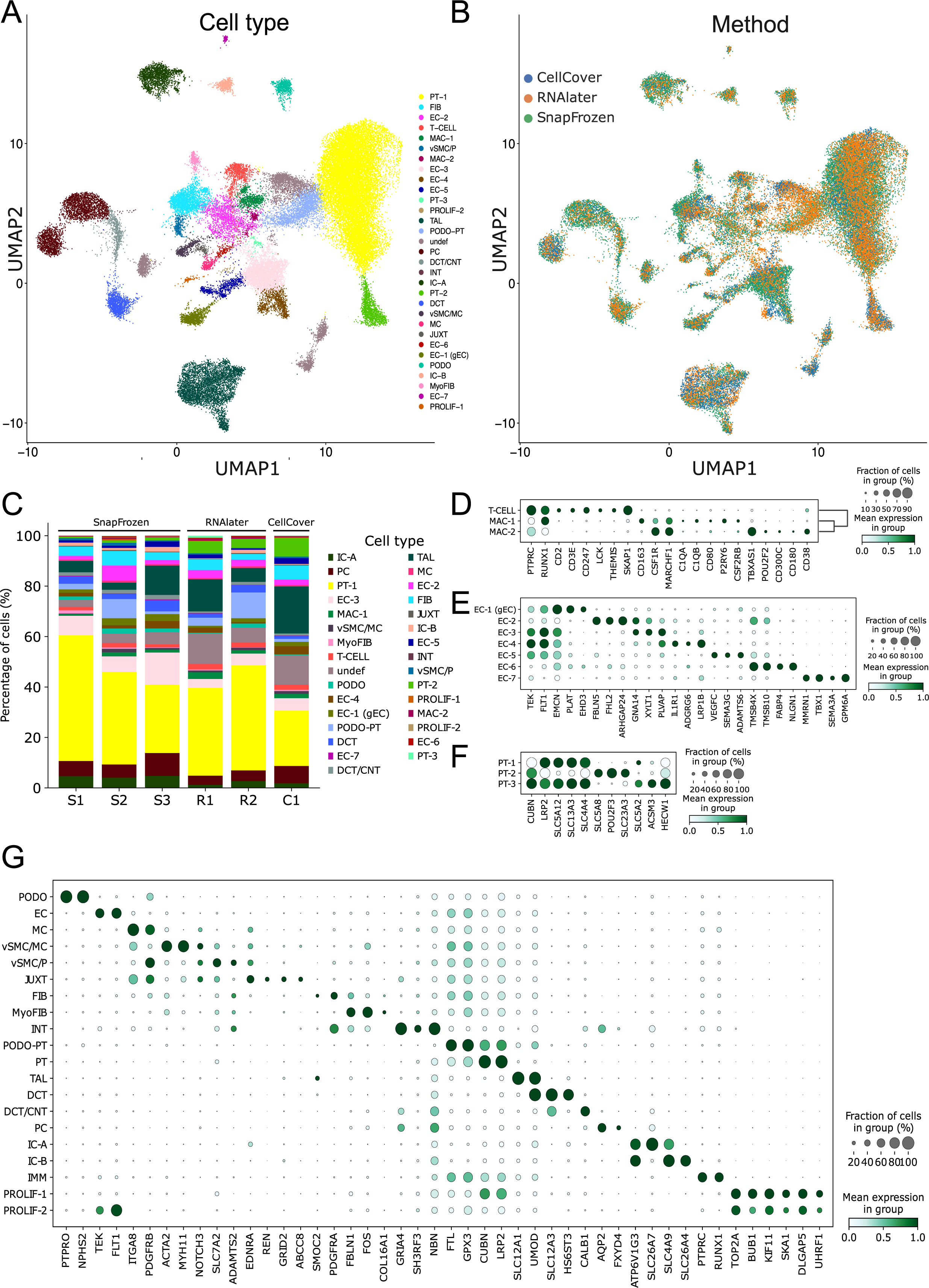
UMAP visualization of a total of 53.351 nuclei from a total of 6 pig kidneys after quality control filtering. UMAP visualization of integrated snRNAseq data sets displays (A) representation of each storage condition in distinct clusters and (B) cell type diversity, each cluster is labeled by color. Samples were integrated using harmony and UMAP was computed on the first 20 principal components after highly variable gene selection. (C) Bar plots showing distribution of cell types from 3 snap frozen (S1-3) and 2 RNA*later* (R1-2) and CellCover (CC) stored kidney biopsies. Dot plot of select average gene expression values (log scale) and percentage of nuclei expressing these genes within each cluster for known and newly discovered cell-type marker genes of (D) major cell types of the kidney as well as of (E) 3 subtypes of proximal tubule cells, (F) 7 subtypes of endothelial cells (EC 1-7) and of (FG 3 subtypes of immune cells (IMM) including t-cells (T-CELL) and 2 types of monocytes/macrophages (MAC-1, MAC-2).

A possible impact of RNA*later* and CellCover on the transcriptome of cryopreserved kidneys was intensively evaluated. The mean expressions of the distinct kidney cell types obtained from tissue stored in RNA*later* and CellCover correlated positively with the gold standard snap frozen (R²=0.82 and 0.75, Figure 4A). To gain further insights into cell-specific preservation-related artifacts, we compared the differential gene expression of each cell-type of RNA*later* or CellCover stored kidney tissue separately with that of snap frozen tissue. In RNA*later* and CellCover stored samples, 24 and 24 cell types, respectively, showed significant upregulation, while 25 and 29 cell types, respectively, had a down regulated gene expression (p < 0.05) when compared to snRNAseq data sets from snap frozen kidney tissue (Figure 4B).

**Fig. 4:**
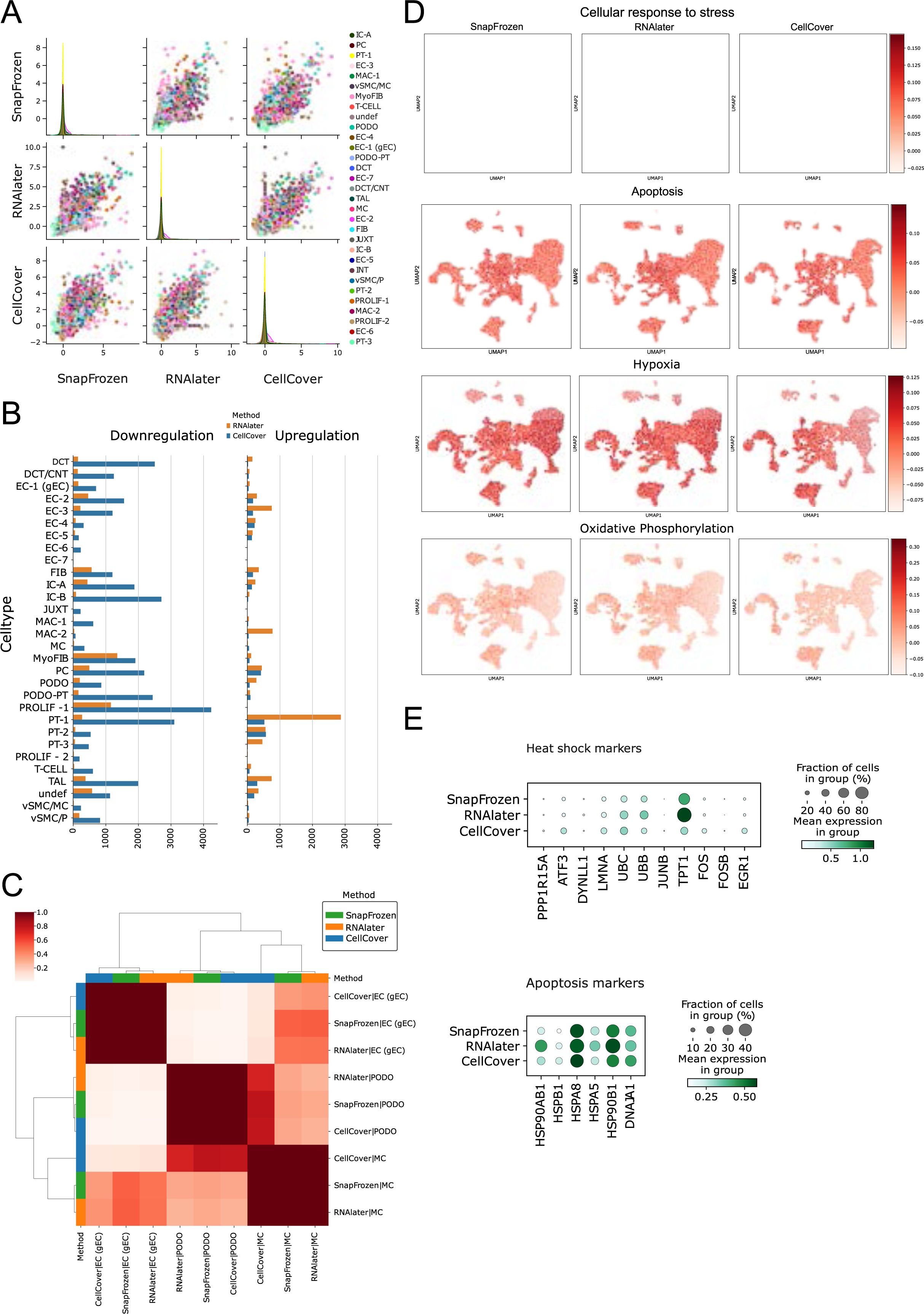
Comparison of cell type specific differential gene expression and cellular stress response between sn RNAseq data sets of snap frozen, RNA*later* and CellCover stored pig kidney tissue. (A) Cell type specific correlation of RNAlater and CellCover stored kidney biopsies with snap frozen kidney biopsies. (B) Cell type specific differential up- and downregulated genes displayed on the x-axis for RNAlater (orange) and CellCover (blue) stored kidney tissue compared to snap frozen kidney tissue. Levels are calculated by Wilcoxon test (as implemented in Scanpy) with logFC = 1.5, FDR < 0.05 as thresholds. (C) Replicability of glomerular cell types between snRNAseq data sets obtained from the different cryopreservation conditions as defined by the area under the receiver operator characteristic curve (AUROC) score. Podo: podocytes, gEC: glomerular endothelial cells, MC: mesangial cells. (D) Comparison of activation of cellular stress response, apoptosis, hypoxia, and oxidative phosphorylation between snRNAseq data sets from snap frozen, RNAlater and CellCover stored kidney tissue. Activation status of each pathway was calculated with AUCell. (E) Violin plots showing the UMI counts for canonical stress related genes in snRNAseq data sets from snap frozen, RNAlater and CellCover stored kidney tissue

Glomerular cells like podocytes, glomerular endothelial cells and mesangial cells represent rare kidney cell types, which are often enriched for scientific purposes as primary cell cultures or from abundant experimental tissue. Transcriptional changes that might be induced by RNA*later* or CellCover would significantly reduce data quality. To rule out transcriptional differences of glomerular cells between single nucleus data sets obtained from kidneys stored in RNAl*ater*, CellCover or snap frozen, we calculated the receiver operator characteristic curve (AUROC) score. using MetaNeighbor [26]. Each glomerular cell type obtained from the respective storage condition had a very high AUROC score for the corresponding cell type obtained from one of the other two storage conditions (Figure 4C), indicating high similarity of cell types obtained by the different storage conditions.

Snap frozen kidney needle biopsies are immediately frozen after collection, allowing less time for the induction of transcriptional changes due to alterations in cellular homeostasis. RNA*later* and CellCover facilitate the handling of kidney biopsies because they can be frozen later. To investigate whether delayed freezing induces cellular stress responses, response to hypoxia or apoptosis, we conducted an enrichment analysis of these pathways with AUCell [27].

In all cell types of the kidney biopsies stored in RNAlater, CellCover or snap frozen we detected a similar activation of the stress response pathways (Figure 4D). Additionally, we found no differences in the expression of genes related to the hypoxia and apoptosis signaling in most cell types under any of the storage conditions tested (Figure 4D). The expression levels of the canonical stress response genes *HSP90AB1*, *HSPB1*, *HSPA8*, *HSPA5*, *HSP90B1* and *DNAJA1* were moderate, and we observed low expression levels of the apoptosis markers *PPP1R15A*, *ATF3*, *DYNLL1*, *LER3*, *LMNA*, *UBC*, *UBB*, *JUNB*, *TPT1*, *FOS*, *FOSB* and *EGR1* in all storage conditions (Figure 4E).

Overall, the single nuclear transcriptome of RNAlater, CellCover or snap frozen stored kidney tissue did not substantially differ, but kidney tissue stored in RNA*later* showed slightly smaller differences in quality control metrics than tissue stored in CellCover compared to snap frozen tissue.

### Single nucleus analysis of healthy and diseased human kidney biopsies

To confirm that porcine kidney tissue constitutes a suitable surrogate for human kidney tissue, fractions from a human kidney allograft biopsy and healthy kidney tissue of 2 tumor-nephrectomies stored in RNA*later* at −80°C were subjected to the snRNAseq pipeline we established with pig kidney tissue. After sequencing and processing the data sets with CellRanger software, the analysis revealed an average estimated number of 15944 nuclei per tumor-nephrectomy sample and a median number of 943.5 genes per nucleus were detected in the tumor-nephrectomy samples. The data set obtained from the diseased kidney biopsy yielded an estimated number of 11313 nuclei and a median number of 2057 genes per nucleus (Supplementary Table 3). After filtering with the described parameter, dimensionality reduction and integration of the 3 data sets with Harmony [15] (Figure 5A), a total number of 43010 nuclei clustered into the diverse cell types of the kidney (Figure 5B) that were annotated on the basis of the expression patterns of cell-specific marker genes reported for previous kidney single-cell data sets (Figure 5C) [10], [28].

**Fig. 5:**
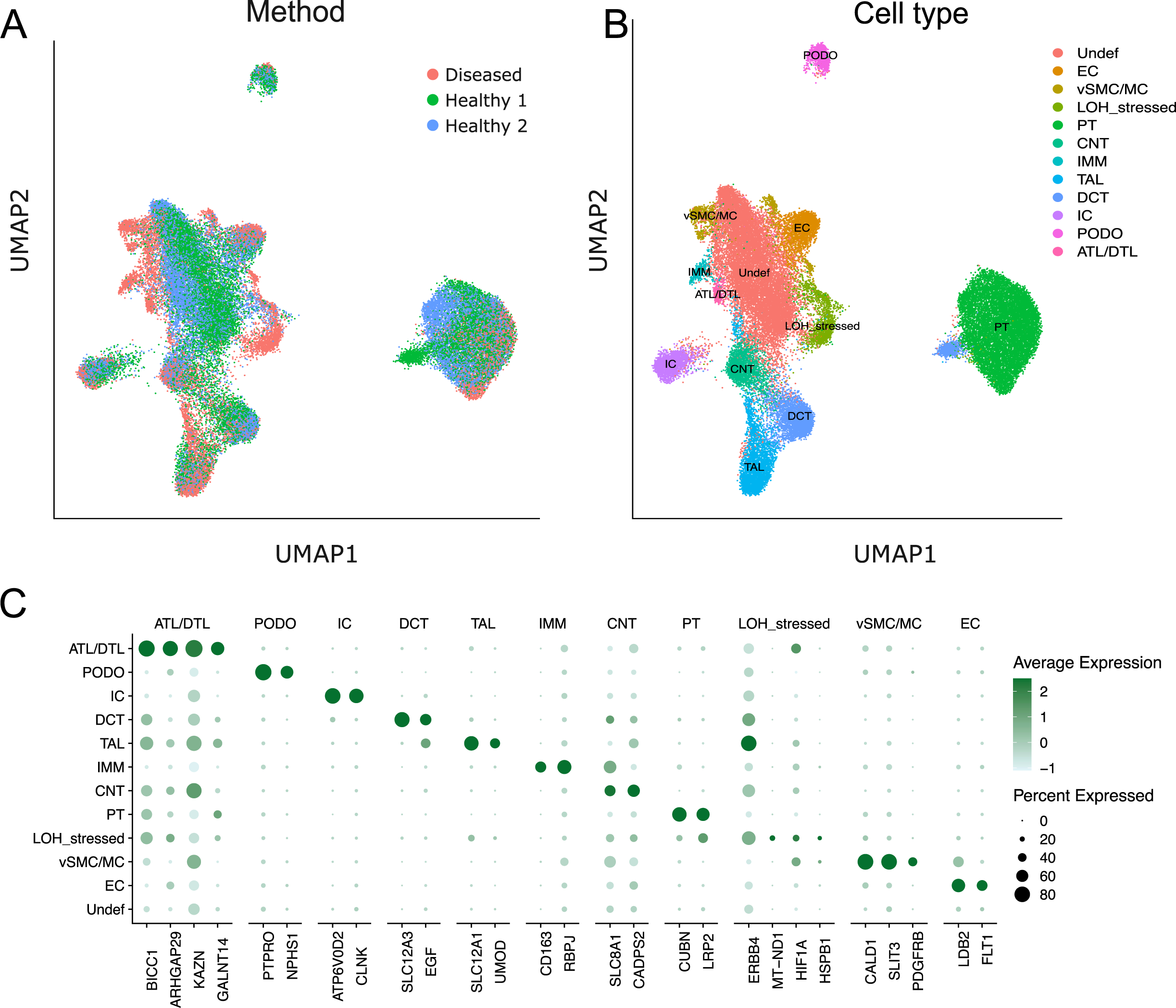
UMAP visualization of 3 snRNAseq data sets from human kidney tissue stored in RNA*later*, (A) after integration of 2 snRNAseq data sets from healthy controls and 1 data set from a clinically indicated transplant kidney biopsy, (B) clustering into 12 distinct cell types of all major cell types of the kidney. (C) Dot plot of cell-type defining marker genes.

### Proteomics of cryopreserved pig kidney tissue

Next, we sought to determine the suitability of RNA*later* and CellCover stored kidney tissue for proteome-analysis by using mass spectrometry. The proteome-analysis of snap frozen, RNA*later* and CellCover stored pig kidney tissue resulted in a complete data set of 1670 proteins (Supplementary Table 4). Globally, the detected protein LFQ intensity across conditions and replicates was consistent (Figure 6A).

**Fig. 6:**
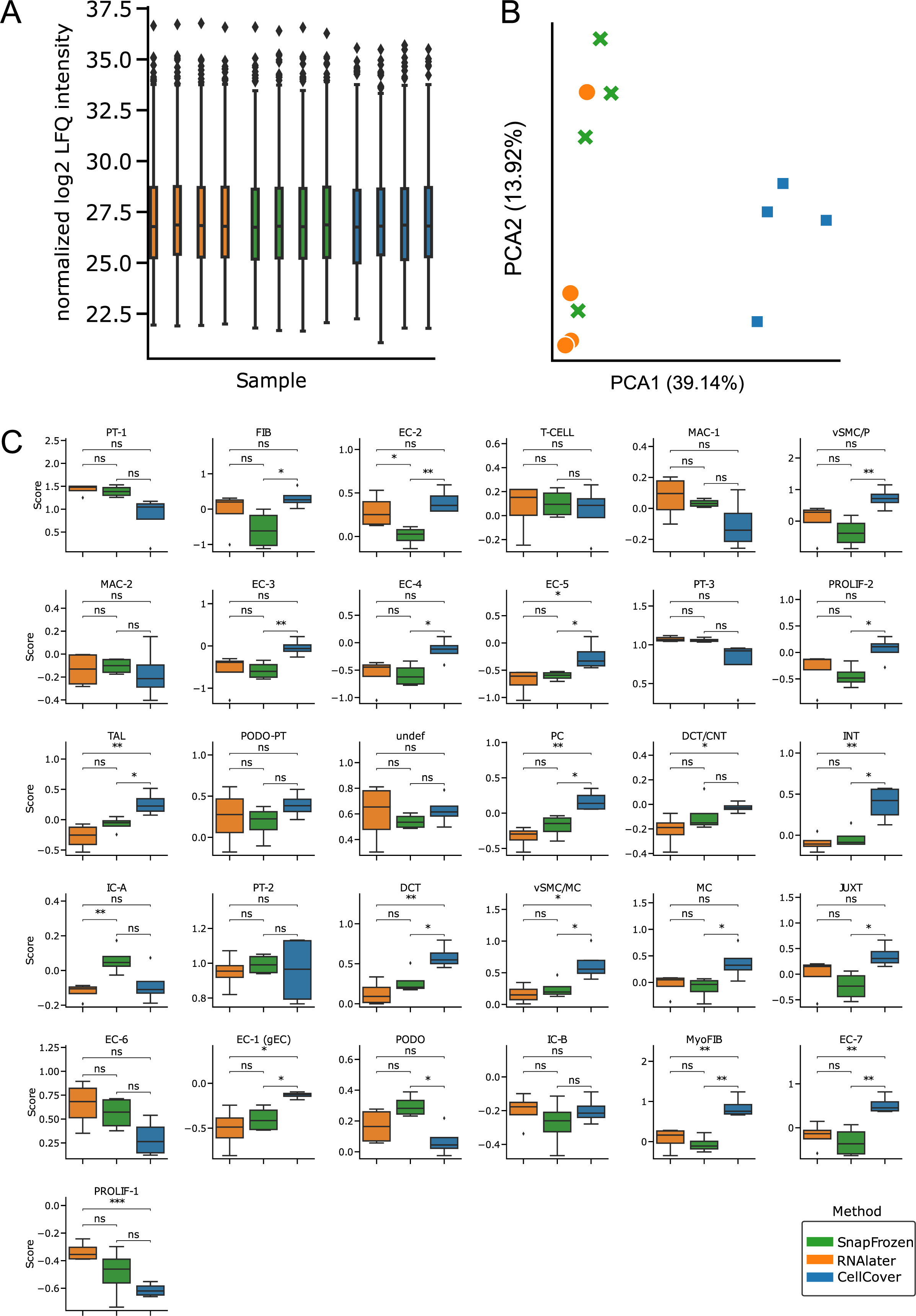
Proteome analysis of 4 kidney biopsies. (A) Distribution of protein abundances (LFQ normalized) of each sample across the three storage methods. (B) PCA representation of the 12 samples across the three methods. (C) Scores of cell type marker genes in proteomics data for the cell types detected in snRNAseq data. The scores were calculated using the score_genes function in Scanpy using the 20 most significant genes with a positive log2FoldChange of 0.1 between cell types as cell type markers. Significance calculations between methods were done using a two-sided t-test.

Principal component analysis between the different cryopreservation methods showed high similarity between tissue stored in RNA*later* compared to snap frozen tissue (Figure 6B), with principal component (PC) 1 being very similar and PC2 playing a smaller contribution to the observed difference (13,92% PC2 vs 39,14% PC1). The proteome of CellCover stored kidney tissue was markedly different along PC1 and somewhat different along PC2 compared to snap frozen tissue. In contrast to RNA*later*, our proteomic analysis indicates that 659 of 1670 proteins had a different abundance when comparing CellCover and snap frozen tissue (adjusted p-value <0.05, Supplementary Table 4).

We further aimed to investigate possible cell type specific differences between RNA*later*, CellCover and snap frozen stored kidney tissue in the proteome. The 20 most upregulated genes for each cell type were extracted and used to compute module scores in the proteome. Two cell type specific differences were found in RNA*later* stored kidney tissue compared to snap frozen tissue, whereas 18 of 31 cell types obtained from CellCover stored tissue showed significant differences in the proteome compared to snap frozen tissue (Figure 6C). In this study we can show that proteome analysis of RNA*later* provides comparable results to snap frozen kidney tissue. Proteome analysis of CellCover stored kidney tissue revealed lower concordance to snap frozen tissue, more than 39% of the proteins had different abundances.

### Metabolomics of cryopreserved pig kidney tissue

In metabolomics, LC-MS and GC-MS are two complementary, commonly used techniques. Metabolites must be volatile or otherwise must be derivatized prior to GC-MS analysis, which limits the compounds suitable for this technique. However, GC-MS delivers excellent chromatographic separation, and its ionization mechanism covers a wide range of compound classes. LC-MS is not as limited as GC-MS regarding analyte properties, harbors good chromatographic separation, but its ionization technique is not as universal as in GC-MS. Thus, we used a combination of targeted LC-MS and non-targeted GC-MS to study the influence of cryo-preservation reagents on the renal metabolome.

In our study, data from non-targeted GC-MS analysis could not be used since small molecules introduced by the two cryoconservation reagents strongly suppressed the signals derived from metabolites. These conservation reagents might also interfere with the derivatization reaction required for GC-MS analyses. In fact, the intensities of the internal standards in the RNA*later* samples decreased to approximately 50% of the snap frozen condition and many metabolites present in the snap frozen samples were not detected at all in either conserved condition. Next, the lipophilic extracts were analyzed by targeted lipid profiling. Mass spectrometric analysis was possible, probably since the conservation reagents and buffer salts remained in the aqueous phase. As shown in the PC analysis, snap frozen control tissue was clearly different from both tested conservation methods (Figure 7A). Remarkably, CellCover’s variance was responsible for most observed alterations, as it scattered around PC1. RNA*later* also exhibited a much higher variation than snap frozen tissue. This is reflected in the averages of the coefficients of variation with 0.07 in the quality control sample, 0.20 in the snap frozen tissue samples, 0.51 in RNA*later* and even 0.62 in CellCover.

**Fig. 7:**
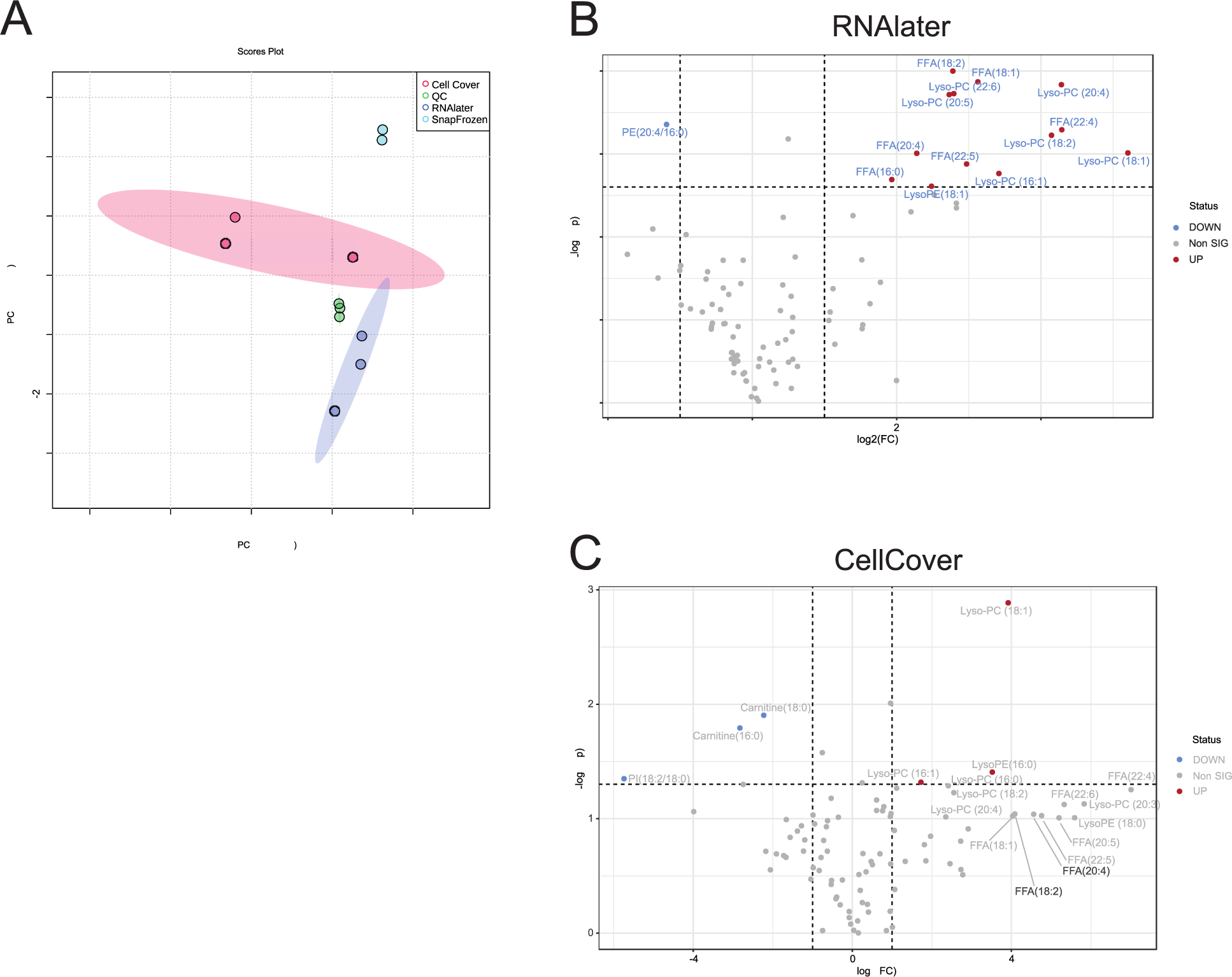
Performance of snap frozen, RNAlater and CellCover stored kidney tissue in metabolome analysis. (A) Principal component analysis of targeted lipid profiling: Each dot represents a sample with their corresponding 95%-confidence interval as shaded area. Each storing condition were different from each other. CellCover and RNA*later* harbored a higher variance than snap frozen and quality control samples. Red: CellCover (n=3), blue: RNA*later* (n=3), turquoise: snap frozen (n=2), green: quality control samples (n=3). Volcano plots of RNA*later* (B) and CellCover (C) compared to snap frozen kidney biopsies. On the x-axis, the log_2_-foldchanges are displayed, while negative log_10_-p-values are displayed on the y-axis. Dashed lines represent the cut-offs of 0.05 for p-value and 2 for fold change. Lipids matching these cut-offs are labeled in blue if down regulated and in red when increased compared to snap frozen. Lipids labeled in grey (b) had a fold change higher than 4 but a p-value between 0.05 and 0.1.

A comparison of lipids altered between RNA*later* and snap frozen revealed several enriched free fatty acids and lyso-phospholipids (Figure 7B). A very similar pattern was observed in samples preserved in Cell Cover (Figure 7C). In the latter comparison, several lipids were enriched more than fourfold compared to snap frozen, but only had a p-value < 0.1. All these lipids were again free fatty acids and lyso-phospholipids. The p-values might be explained by the high variance in this group. A specific accumulation of these two lipid classes indicates an insufficient metabolic quenching by still active lipases. While snap frozen kidney tissue is suitable for metabolomics with current technologies, RNA*later* and CellCover stored kidney are not.

### Histopathological evaluation of tissue architecture and antigenicity of cryopreserved pig kidney tissue

To address whether RNA*later* and CellCover preserve kidney tissue structural architecture and molecular antigenicity, conventional histopathological with periodic acid-Schiff (PAS)and immunofluorescence (IF-) staining from pig kidney stored in RNA*later* and CellCover were performed and compared to OCT-embedded tissue as control.

To determine the optimal storage conditions, pig kidney tissue stored in RNA*later* and CellCover were either placed immediately in −20°C after an initial short incubation at 4°C for 30 to 60 minutes or underwent an automated, controlled cool-down at a rate of −1°C/minute to −20°C. RNA*later* embedded kidney tissue was well preserved at −20°C, independent of immediate (Figure 8A) or automated, controlled (Supplementary Figure 2) freezing condition. CellCover preserved kidney tissue appeared to be less well preserved compared to RNA*later* embedded tissue, as a higher proportion of cells were detached from the basement membrane (Figure 8B). No difference in IF-staining signal quality was observed between RNA*later* and CellCover preserved kidney tissues (Figure 8C and D). Structural architecture and molecular antigenicity of RNA*later* and CellCover preserved kidney tissue was additionally examined after freezing from −20°C to −80°C for long time storage.

**Fig. 8:**
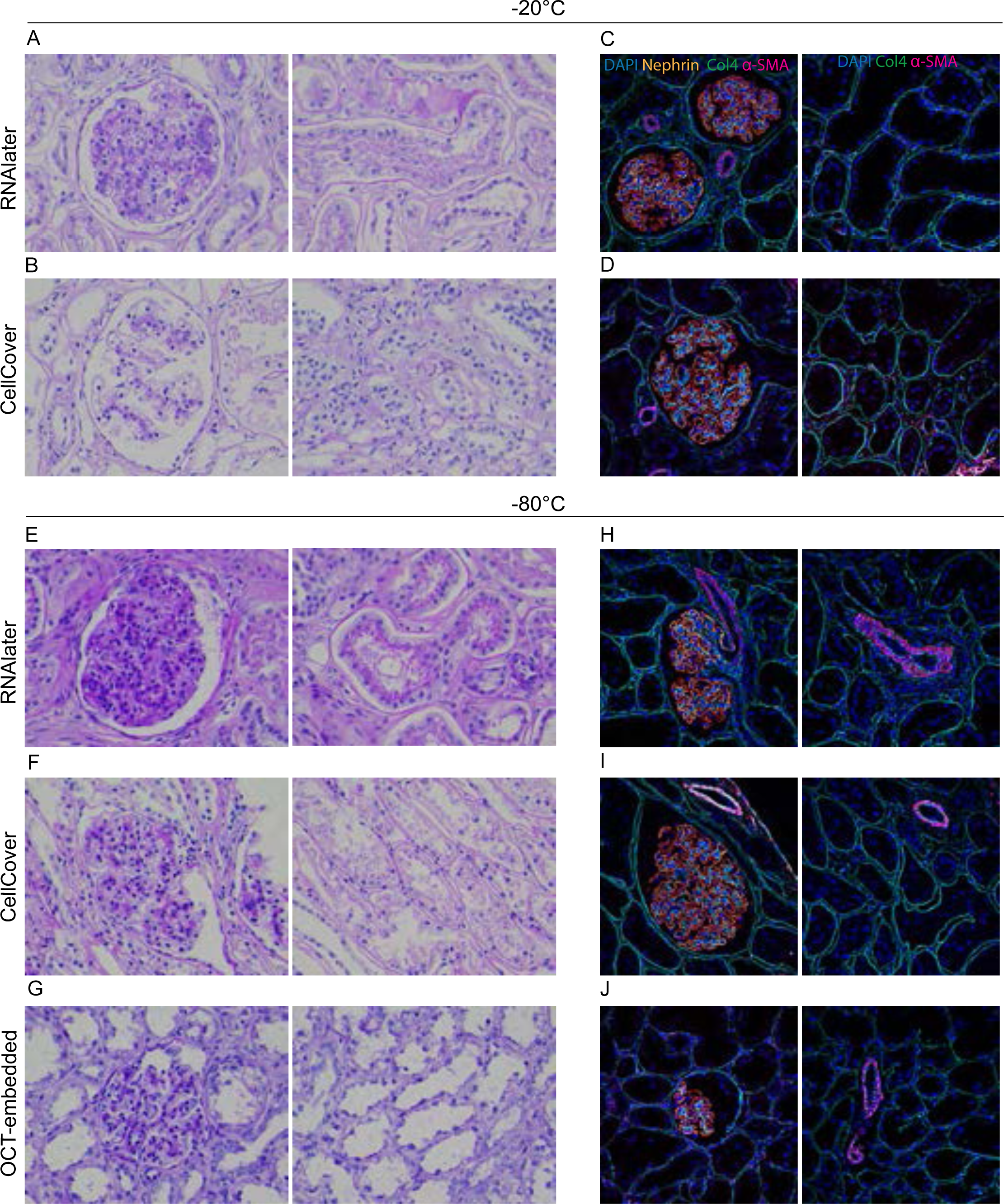
Assessment of tissue architecture preservation (PAS-staining) and immunofluorescence staining of RNAlater and CellCover stored kidney tissue compared to OCT-embedded tissue. Immunofluorescence staining of nuclei (DAPI), basement membrane (Collagen 4), slit diaphragm (Nephrin) and smooth muscle cells (α-SMA) of pig kidney tissue. PAS-Staining of RNAlater (A) and CellCover (B) stored kidney tissue immediately frozen to −20°C. PAS-Staining of RNAlater (C) and CellCover (D) stored kidney tissue at −80°C and OCT-embedded kidney tissue (E). Immunofluorescence staining of RNAlater (F) and CellCover (G) stored kidney tissue immediately frozen to −20°C and RNAlater (H) and CellCover (I) stored kidney tissue at −80°C and OCT-embedded kidney tissue (J).

Freezing of RNA*later* and CellCover preserved kidney tissue from −20°C to −80°C did not introduce more freezing-related artifacts (Figure 8E and F) as freezing to −20°C only or as OCT-embedding (Figure 8G). Again, we found that kidney tissue frozen in CellCover appeared to be less well preserved compared to RNA*later* preserved tissue. Similarly, molecular antigenicity was preserved in both RNA*later* and CellCover embedded kidney tissues that underwent freezing from −20°C to −80°C as shown by IF-staining (Figure 8H and I) and yielded comparable results to OCT-embedded kidney control-tissue (Figure 8J).

Overall, this study could demonstrate that both RNA*later* and CellCover preserve kidney tissue molecular antigenicity at −80°C. However, the structural architecture appeared to be more intact in RNAlater stored tissue.

## Discussion

Single cell transcriptomic studies contributed to substantial progress in understanding kidney tissue heterogeneity and provided greater insight into the complex cell type composition and orchestrated signaling pathways in healthy and diseased tissue [10], [11], [23]. This technique opens doorways for the identification of targeted therapy and prognostic biomarkers. However, kidney tissue stored in biobanks is cryopreserved and therefore, depending on the preservation method, usually not suitable for scRNAseq since these methods require intact viable single cells. DMSO-containing cryopreservation media, which maintain cell viability during freezing, storage and thawing require liquid nitrogen for storage [30], [31], that mostly limits storage capacities of biobanks. Published single nucleus dissociation protocols have demonstrated substantial advantages including greater flexibility with time and place of analysis, the representation of rare and fragile kidney cell types, reduction of dissociation induced transcriptional stress responses and comparable sensitivity to scRNAseq [10], [23]. These findings are crucial for ushering in a new era of translational research, as single cell analyses are no longer limited to mainly experimental tissue. One hurdle still to be overcome is the collection of high-quality kidney tissue for biobanks, which should be translatable into clinical practice. Although comprehensive studies examined the advantages of single nucleus over single cell dissociation and highlighted the compatibility of archived frozen tissue [23], [32], [33], an evaluation of a potential kidney cell-type specific transcriptional impact of preservation media such as RNA*later* or CellCover, which facilitate tissue collection for repositories in daily clinical practice, has been lacking. Thus, we optimized a tissue processing pipeline for kidney biopsies stored in RNA*later*, that allows snRNAseq, proteome and histopathological analysis.

First, we demonstrated that our adapted robust single nucleus dissociation protocol for 3-4 mm of snap frozen or RNA*later* and CellCover stored kidney biopsy cores yielded in sufficient numbers of intact nuclei for further processing with the high throughput single cell platform Chromium 10X Genomics. High throughput technologies permit the analysis of thousands of cells in parallel if enough high-quality single cells/nuclei can be provided. Especially when investigating less abundant cell populations like glomerular cells, the total number of profiled cells is crucial [34].

Sequencing quality control metrics revealed significantly lower fractions of ambient RNA but higher mitochondrial contamination in snRNAseq data sets obtained from RNA*later* stored kidney tissue compared to snap frozen tissue. In snRNAseq data sets from CellCover stored tissue, both mitochondrial and ambient RNA contamination was found to be significantly higher as in snap frozen snRNAseq data sets. Although mitochondrial transcripts are not expressed in the nucleus, they were reported to be present in variable quantities in snRNAseq data sets due to association with the nuclear membrane [10]. Elevated mitochondrial transcript are indicators for cell stress [35], but enrichment analysis of the single nuclear transcriptome of snap frozen, RNA*later* and CellCover stored kidney tissue did not uncover relevant differences in apoptosis, hypoxia and cellular response to stress pathways. Two possible causes can underlie these controversial findings. First, RN*Alater* and CellCover could increase mitochondrial adhesion to the nuclear membrane. The preservation media might alter membrane surfaces which could also explain the finding of slightly increased multiplets in RNA*later* and CellCover preserved samples compared to snap frozen. Second, each data set was generated from an individual (sacrificed) pig with possibly interindividual varying mitochondrial abundance. However, our data sets show a low overall mitochondrial transcripts abundance of < 2.5% and do not exceed previous reported numbers of mitochondrial contamination [10], [9], [10].

Applying all filter criteria to the snRNAseq data sets, the number of recovered nuclei exceeded previously published results for snap frozen [36] and RNA*later* stored kidney tissue [9]. Depending on the underlying kidney disease, cell type specific transcriptional changes induced by RNA*later* or CellCover might lead to misinterpretation of results for disease mechanisms, potential targets, and biomarkers. Thus, we performed cell type specific differential expression analysis between RNAl*ater* or CellCover and snap frozen stored kidney tissue. Up-regulation of genes was overall low in both preservation methods, CellCover showed somewhat more downregulated genes over all cell types. This finding might be attributed to preservation medium specific transcriptional changes or interindividual differences of the respective pig kidneys.

As fractions of kidney biopsies provide limited numbers of glomeruli and glomerular cells represent lower proportions of the kidney [34] transcriptional programs altered by RNA*later* or CellCover will have higher impact as in abundant cell types. We could not detect any influence of the storage condition of the single nuclear transcriptome of 3 glomerular cell types (podocytes, glomerular endothelial cells, mesangial cells).

In contrast to previous studies, which commonly reported a lower detection rate of immune cells in snRNAseq data sets compared to scRNAseq data sets [37], [10], [23], [38] our protocol was able to recover a total of 1896 immune cells in the data sets of pig kidney, clustering into t-cells and monocytes/macrophages. To minimize the loss of rare cell types such as immune cells, the entire protocol was performed on ice, the number of sieving and centrifugation steps was reduced, and a harsh lysis buffer shortened the dissociation time by rapidly releasing even strong cellular associations. Incomplete cell dissociation in scRNAseq protocols might lead to under-representation of resident cell types and artificially increases leukocyte rates in the tissue [38].

We further demonstrated that porcine kidney tissue represents a surrogate for human kidney tissue, as we were able to apply our protocol without any modification to 1 human diseased kidney biopsy and 2 healthy controls (tumor-free tissue from tumor-nephrectomies). These findings are consistent with previously published results for heart tissue [33].

Overall, our rigorous snRNAseq data analysis of RNA*later* stored kidney tissue could outline that RNA*later* constitutes an adequate alternative for immediate tissue preservation if snap freezing of tissue is not implementable into clinical routines. In addition to slightly higher concordance of quality control metrics of snRNAseq data sets from RNAlater and snap frozen kidney tissue, a major advantage of RNAlater over CellCover is its long-time experience in tissue RNA protection [6], [39].

As our optimized protocol requires only 3-4 mm of a biopsy core for high throughput snRNAseq, the remaining part of a biopsy core stored in RNA*later* or CellCover can be utilized to generate other omics data. Tissue stored in RNA*later* can be refrozen in contrast to CellCover and snap frozen tissue, therefore further analysis techniques are independent of time and place. Although samples stored in RNA*later* have been used extensively in DNA and RNA studies, only a few studies have investigated the feasibility in retrieving proteins, metabolites or histopathological information from human tissue samples stored in RNA*later* [40], [41], [2], [3].

The investigation of one kidney biopsy core stored in RNA*later* with additional data dimensions by combining single cell resolution of transcriptome data with bulk data of intra- and extracellular proteins as well as spatial resolution of imaging [42] will be a major step in translational medicine and overcome the need of disease model systems. In addition, the generation of these data sets provides the basis for the need to develop novel bioinformatics and mathematical modeling tools and represents a further step towards personalized medicine, as Riedel et al. were already able to demonstrate glucocorticoid response in patients with crescentic glomerulonephritis by analyzing histological and single cell data obtained directly from fresh kidney biopsies [43]. As previously reported for tumor tissue of pancreatic ductal adenocarcinoma [41], we could demonstrate that mass spectrometry proteome analysis of RNA*later* stored kidney tissue matched highly with snap frozen kidney tissue and no protein showed a significant difference between both storage conditions.

In addition, we computationally generated cell type-specific modules with sn RNAseq data sets and compared these modules in the proteomic data between the three storing conditions. The cell type specific modules from RNA*later* stored kidney showed comparable protein abundance to those from snap frozen tissue. More differences were detected in CellCover stored kidney tissue. For proteome analysis, only one pig kidney was preserved by the different preservation methods, thus the differences found for CellCover stored tissue can be attributed to the medium.

The metabolome is relevant for understanding of physiological and disease-modifying mechanisms as well as its interaction with the transcriptome and proteome of the kidney is only beginning to be understood [44]. However, this study illustrates that with current technologies only snap frozen kidney biopsies can be used for metabolomics studies. Preservation of kidney tissue architecture is essential for bio banked tissue, as morphologic examination of kidney tissue biopsies is still essential for histopathological diagnosis. In addition, preserved architecture of archived tissue might constitute added value not only for scientific efforts, but also for ethical aspects of kidney tissue biobanks. Kidney tissue stored in RNA*later* could be used as supplemental material if the kidney tissue collected for clinical histopathology does not contain enough informative material and would otherwise require a new invasive procedure for a definitive diagnosis of the underlying kidney disease.

Moreover, disease-specific morphological and molecular alterations can be assessed by artificial intelligence (AI) -driven pattern extraction and improve disease classifications, pathogenesis and prognosis as well as therapy response [45], [46]. AI-defined morphological and molecular patterns can be integrated with data obtained from other, complementary omics like sn transcriptomics and proteomics. In our study we show that RNA*later* mostly preserves kidney architecture and does not interfere with immunofluorescence staining. We tested PAS and a limited number of antibodies for immunofluorescence staining of RNA*later* preserved porcine kidney tissue. In contrast to our study, a previous study demonstrated impaired histological structure and immunohistochemical staining patterns for mouse kidneys stored in RNA*later* at −20°C. These contrasting results may be explained by a longer incubation time of the mouse tissue at 4°C overnight prior to freezing to −20°C [47]. CellCover stored kidney tissue was less suitable for histopathological examination in our hands.

However, our study has several limitations that require further investigation. First, additional studies with a larger number of antigens and with different types of imaging on human tissues are needed to further evaluate the preservation of tissue morphology and the protein-preserving effect of *RNA*later. Second, we did not investigate whether or how the duration of tissue cryopreservation and long-term storage affects the dissociation and integrity of cell nuclei, the single-cell transcriptome landscape or proteome, and the structural architecture and molecular antigenicity of kidney tissue. Finally, we did not examine the transcriptomic landscape of single cells between fresh and cryopreserved tissue preparations.

In conclusion, our study shows that RNA*later* can facilitate the collection and storage of kidney tissue by bypassing the need for snap freezing in liquid nitrogen without compromising snRNAseq, proteome and histopathological analysis. Only metabolome analysis is currently limited to snap frozen stored kidney biopsies.

This work will guide future efforts to illuminate the molecular landscape of kidney diseases by applying multi-omic approaches to translational medicine, enabled by the integration of tissue collection for biobanking into clinical routines.

## Authors Contributions

S.E.G., S.H. and M.T.L. conceptualized, designed, conducted, and interpreted experiments and analyses. R.K., F.H. and S.H. designed and conducted the bioinformatic analyses. M. P. and T.Z. assisted with data handling. D.K, S.L., M.La., B.K. conceived and analyzed experiments. S.Lu and S.Liu provided input in experimental design and assisted with data interpretation. S.C. and I.E. performed experiments. J.C. provided critical samples.

F.B., V.G.P., M.M.R., T.B.H and S.B. gave input in experimental design and provided critical review and commentary. S.E.G, S.H., M.T.L. and T.B.H. supervised the project. S.E.G., S.H., M.T.L, and T.B.H. wrote the manuscript with input and discussion from all authors.

## Acknowledgements

The authors acknowledge excellent technical support by Stefan Gatzemeier. R.K. was supported by FOR5068 P9 and the 3R initiative of the UKE; F.H. by the M3I excellence initiative and a UKE postdoctoral stipend; and S.B. by SFB1286 SP02, SFB1192 B8, and C3. S.H. was supported by SFB1192 B8. F.B. was supported by the Else Kröner-Fresenius-Stiftung (2021_EKMS.26) and the German Society of Nephrology. D.K. received intramural funding (Clinician/Scientist program). M.M.R was supported by the DFG (RI 2811/1-1 and RI 2811/2-1, FOR2743, and SFB1192-project B10), by the Young Investigator Award from the Novo Nordisk Foundation, grant number NNF19OC0056043, the Carlsberg Young Investigator fellowship as well as Aarhus universitet forskningsfond, and an endowed professorship by the Klinikum Bad Bramstedt. M.M.R. thanks the Klinikum Bad Bramstedt for support. This project was also supported by the European Union’s Horizon 2020 research and innovation funding programme under the Marie Skłodowska-Curie grant agreement No 754513 and The Aarhus University Research Foundation (to M.M.R). M.T.L was supported by BMBF, grant 01GM1518A; STOP-FSGS and BMBF grant 01EK2105D, UPTAKE. T.B.H was supported by the DFG (CRC1192, HU 1016/8–2, HU 1016/ 12–1), by the BMBF (STOP-FSGS-01GM1901C, UPTAKE 01EK2105D). This project has received funding from Abbvie and the Innovative Medicines Initiative 2 Joint undertaking under grant agreement No. 115974 (Beat-DKD) (to T.B.H). This Joint undertaking receives support from the European Union’s Horizon 2020 research and innovation program and EFPIA with JDRF. This work was partially funded by the Else Kröner-Fresenius-Stiftung and the Eva Luise und Horst Köhler Stiftung – Project No: 2019_KollegSE.04 (to S.G. and T.B.H).

