## Supplementary figures and images for "A universal preservation protocol for multi-omic and histological analysis of kidney tissue"

### Suppl Fig. 1

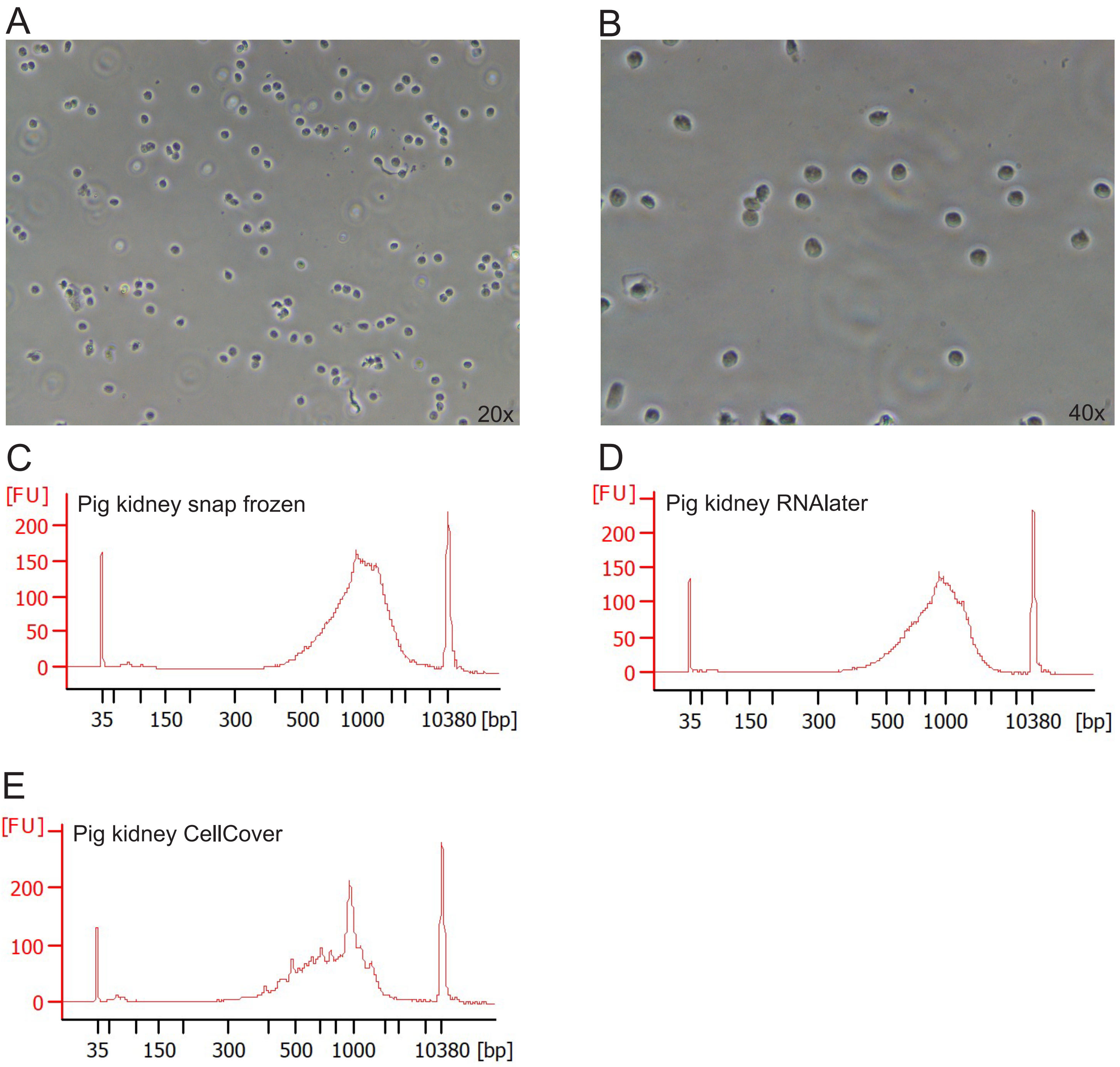

### Suppl Fig. 2

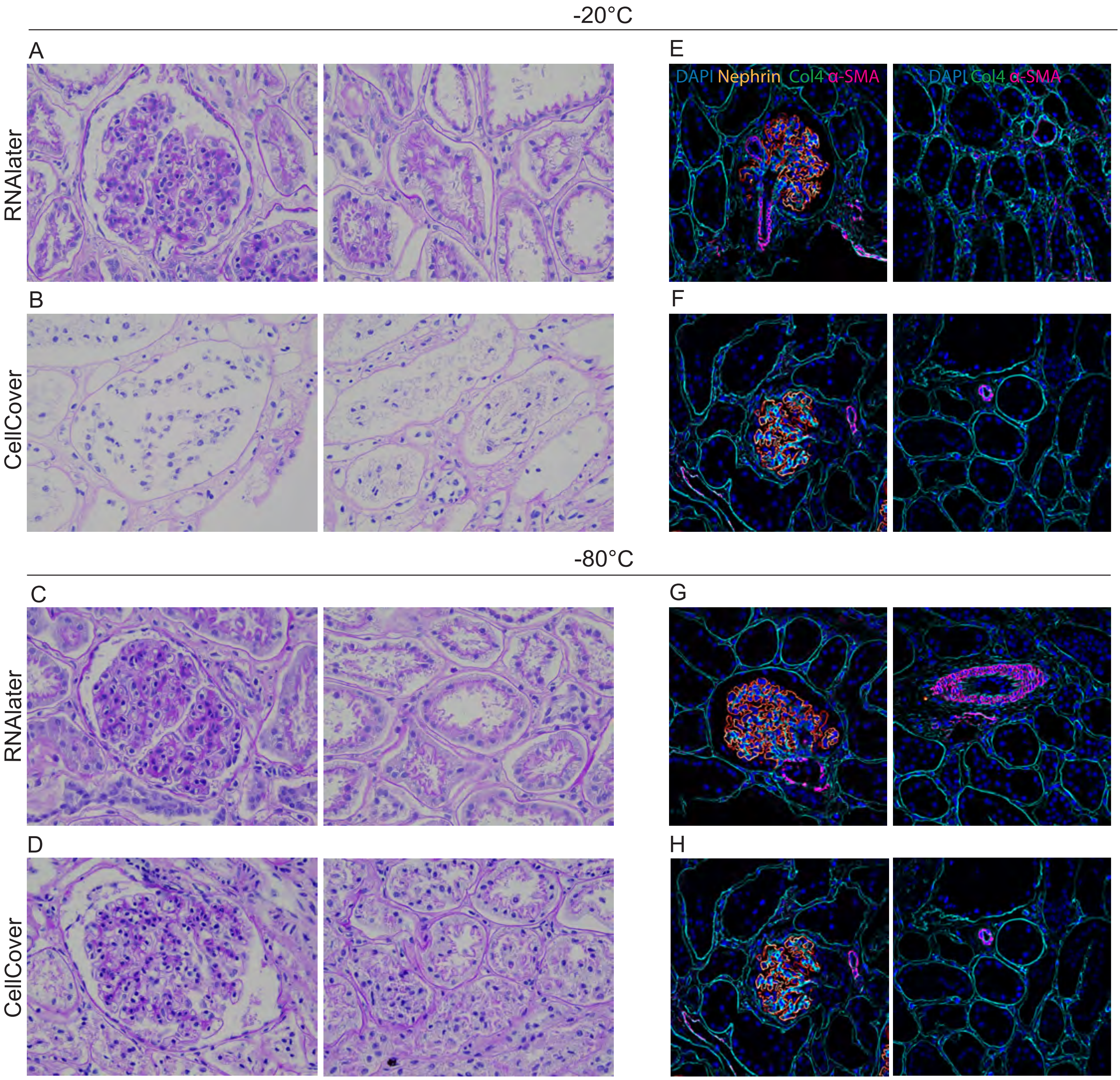
