## Supplementary material for "A universal preservation protocol for multi-omic and histological analysis of kidney tissue": Suppl Tab. 1

| Target | Manufacturer | Catalog No. | Species | Dilution |
|---|---|---|---|---|
| aSMA/FITC conjugate | Abcam | F3777 | Mouse | 1:200 |
| Collagen IV | Abcam | Ab6586 | Rabbit | 1:200 |
| Nephrin | Progen | GP-N2 | Guinea pig | 1:100 |
